# Before auxin distribution matters: Prepatterning of the Arabidopsis embryo axis by MPK6-dependent AUX/IAA8 phosphorylation

**DOI:** 10.64898/2026.08.09.743751

**Authors:** Torren Bischoff, Marina Ortega-Perez, Alexa-Maria Wangler, Helen Schäfer, Martina Kolb, Kai Wang, Houming Chen, Yingjing Miao, Yashodar Babu, Lisa Asseck, Laura Ragni, Daniel Slane, Gerd Jürgens, Martin Bayer

## Abstract

In growth and development of plants, the signaling molecule auxin plays an instrumental role. Many patterning processes are controlled by auxin signaling in a self-organizing matter. After fertilization, such a patterning process needs to be started for the first time during plant life. Contrary to previous models, we show that the first differential auxin response that separates embryonic from extra-embryonic development in the two zygotic daughter cells happens independently of a detectable auxin gradient. Instead, differential phosphorylation of the transcriptional inhibitor Aux/IAA8 by MAP kinase signaling leads to differences in auxin sensitivity between the two daughter cells, initiating embryonic development in the apical cell. IAA8 phosphorylation in the basal cell prevents proteasomal degradation, suppressing canonical auxin responses, which initiates extra-embryonic development. MAP kinase signaling therefore precedes auxin gradients and prepatterns the apical-basal axis independently of an auxin gradient.

## Introduction

The phytohormone auxin plays a central role in plant development and has been attributed with a morphogen-like function, shaping cell identities and growth with distribution gradients on tissue- and organ level by its unique ability to be transported on a cellular and organ level (Abley *et al*., 2016; Benkova *et al*., 2003; Bennett *et al*., 2014; Bhatia and Heisler, 2018; Huang *et al*., 2026; Rutten *et al*., 2022). Several systems perceive auxin concentrations on the cell surface and in the cell (Chu *et al*., 2026; Vanneste *et al*., 2025). The best studied control of auxin-dependent gene expression is exerted by the ubiquitin ligase-based transcriptional auxin response module. Located in the nucleus, this system consists of AUXIN RESPONSE FACTOR (ARF) transcription factors, their highly dynamic INDOLE-3-ACETIC ACID INDUCIBLE (Aux/IAA) interactors, and TIR1/AFB E3 ubiquitin ligases that facilitate Aux/IAA ubiquitination and proteasomal degradation (Dharmasiri *et al*., 2005; Gray *et al*., 2001; Kepinski and Leyser, 2005; Maraschin Fdos *et al*., 2009; Ramos *et al*., 2001). Aux/IAA proteins bind ARFs through their complementary C-terminal PB1 domains (Guilfoyle, 2015; Korasick *et al*., 2014). This complex recruits TOPLESS (TPL)-related proteins which serve as scaffolds for HISTONE DEACETYLASEs (HDACs) and facilitate the suppression of auxin-responsive target genes by chromatin remodeling (Szemenyei *et al*., 2008). At higher auxin concentrations, auxin can serve as molecular glue and enables binding of Aux/IAAs to TIR1/SCF complexes. Subsequent ubiquitination leads to degradation of Aux/IAA proteins by the proteasomal pathway, dramatically lowering the effective Aux/IAA concentration in the nucleus. Low Aux/IAA concentration at permissive auxin concentrations allows ARF activity by the action of HISTONE ACETYLASES, which results in activation of auxin-responsive target genes (Chu *et al*., 2026; Weijers and Wagner, 2016). Since Aux/IAA and ARF genes themselves are in many cases expressed in an auxin-dependent manner, this module allows a bistable output with hysteresis and is believed to be the basis for self-organizing principles in patterning processes (Abley *et al*., 2016; Bhatia and Heisler, 2018; Burian *et al*., 2026; Grieneisen *et al*., 2007; Lau *et al*., 2011).

Recently, another level of TIR1-dependent control of ARF activity has been reported: TIR1 can act as adenylate cyclase to locally produce the second messenger cAMP during the auxin-mediated TIR1-Aux/IAA interaction at the ARF complex. It has been proposed that local cAMP accumulation at permissive auxin concentrations stimulates ARF activity, introducing a second layer of ARF regulation based on the Aux/IAA-TIR1 interaction (Chen *et al*., 2025a; Qi *et al*., 2022). Auxin concentrations in the cell are influenced by local synthesis, degradation, and conjugation, as well as by auxin transport (reviewed in Vanneste *et al*., 2025). The main contributors to directional auxin transport are auxin efflux facilitator proteins of the PIN-FORMED (PIN) family that can be polarly localized at the plasma membrane (Friml *et al*., 2002; Friml *et al*., 2003). Directional auxin transport and auxin response are interdependent and theoretically enable a bistable output with hysteresis that can be the basis for self-organizing principles in patterning processes (Abley *et al*., 2016; Bennett *et al*., 2014; Bhatia and Heisler, 2018; Burian *et al*., 2026; Grieneisen *et al*., 2007; Lau *et al*., 2011).

Auxin signaling plays a decisive role during patterning of the plant embryo. In the *Brassicaceae* family, cell divisions in the early Arabidopsis embryo follow a stereotypical pattern (Johri *et al*., 1992). After fertilization, the zygote elongates and divides asymmetrically to form two daughter cells with different developmental fates (Maheshwari, 1950; Wang *et al*., 2019). The larger basal daughter cell forms the filamentous suspensor, a mostly extra-embryonic support structure, while the smaller apical cell develops into the embryo proper. The embryo undergoes several, tightly controlled rounds of symmetrical and asymmetric divisions forming the precursors of essential plant organs and the stem cell niches (meristems) necessary for post-embryonic development (Dresselhaus and Jurgens, 2021). A hallmark of initiating embryonic development in *Brassicaceae* is the switch from stereotypical horizontal divisions seen in the descendant of the basal cell to a vertical division plane in the apical cell that initiates the 3-dimensional growth of the embryo (Hamann *et al*., 1999; Yoshida *et al*., 2014). This change in division plane orientation as well as many of the following decisive asymmetric cell divisions are directed by auxin signaling. For example, the uppermost cell of the suspensor, termed hypophysis, divides asymmetrically to initiate the formation of the organizing center of the later root apical meristem. This crucial step is controlled by the well characterized ARF -Aux/IAA pair MONOPTEROS (MP) and BODENLOS (BDL/IAA12) (Hamann *et al*., 2002; Weijers *et al*., 2006) and loss-of-function mutations in *MP* as well as gain-of-function mutations in *BDL*that interfere with IAA12 protein turnover lead to rootless seedlings. Expression of *MP* and *BDL* depends on ARF activity itself (Lau *et al*., 2011) and they are already differentially, apically expressed in the two daughter cells of the first zygotic division.

The embryo itself produces auxin only after the 8-cell stage and early development relies on maternally supplied auxin imported from maternal seed tissue at the basal pole of the suspensor (Robert *et al*., 2018). PIN7 is specifically expressed in suspensor cells and localizes to apical plasma membranes. Consequently, PIN7 in the basal cell is believed to export auxin towards the apical cell, establishing a first auxin response that can be detected by a synthetic auxin responsive reporter gene (*DR5rev::erGFP*) (Friml *et al*., 2003). Many *PIN* genes themselves are auxin-inducible, suggesting that differential expression of the MP/BDL-dependent auxin response module in the early embryo is either established by self-organizing principles or follows already existing differences in a polarized embryo.

One molecular pathway that has a well characterized function in polarizing the zygote and establishing differential gene expression in its two daughter cells is the embryonic ERECTA-YODA (ER/YDA) pathway. This Toll-like receptor kinase-MAP kinase pathway comprises receptor kinases of the ERECTA family (ERf) and SOMATIC EMBROGENESIS RECEPTOR-LIKE KINASE family (SERKs), membrane-associated kinase-like proteins of the BRASSINOSTEROID SIGNALING KINASE (BSK) family and the MAP kinase cascade including the MAP3K YODA (YDA), MAPK kinases MKK4 and MKK5, as well as the MAP kinases MPK3 and MPK6 (Wang *et al*., 2019; Wangler *et al*., 2026). This pathway seems to be active in the zygote and its basal daughter cell where it is responsible for differential expression of *WUSCHEL-RELATED HOMEOBOX 8* (*WOX8*) by phosphorylation of the transcription factors INDUCER OF CBF3 EXPRESSION1 (ICE1) and WRKY2 (Chen *et al*., 2025b; Ueda *et al*., 2017; Ueda *et al*., 2011).

Here we report that the first auxin response in the Arabidopsis embryo is independent of a detectable auxin gradient but depends on differential MAP kinase-dependent phosphorylation and stabilization of the Aux/IAA protein IAA8. Differential IAA8 turn-over essentially forms a pre-pattern in the embryo, suppressing ectopic auxin maxima in the basal cell while priming the apical cell for the first auxin response and embryonic development.

## Results

Previous reports showing PIN7 localization at the apical membrane of the basal daughter cell and weak but specific transcriptional activity of the synthetic auxin response reporter gene *DR5rev::erGFP* in the apical cell imply that the first embryonic auxin response is established by an apical-basal auxin gradient (Friml *et al*., 2003). We tested this hypothesis with the ratiometric fluorescence reporter R2D2 that is based on auxin-dependent protein turnover (Liao *et al*., 2015). In conflict with the current model, we could not detect higher auxin-dependent DII-GFP protein turnover in the apical cell (Figure 1A, B) in comparison to the basal cell while we could confirm the previously reported DR5 activity in apical cell (Figure 1C). Only from the 2-to 4-cell stage onwards, we could detect significantly more auxin-dependent protein turnover in the apical embryonic cells than in the suspensor cells by R2D2 (Figure 1A, B), suggesting that an apical-basal auxin gradient is only established after the embryo-specific deviations in cell division patterns have already occurred. The delayed difference in auxin response therefore appears to be a consequence rather than the cause for the initial difference between apical and basal cells.

**Figure 1:**
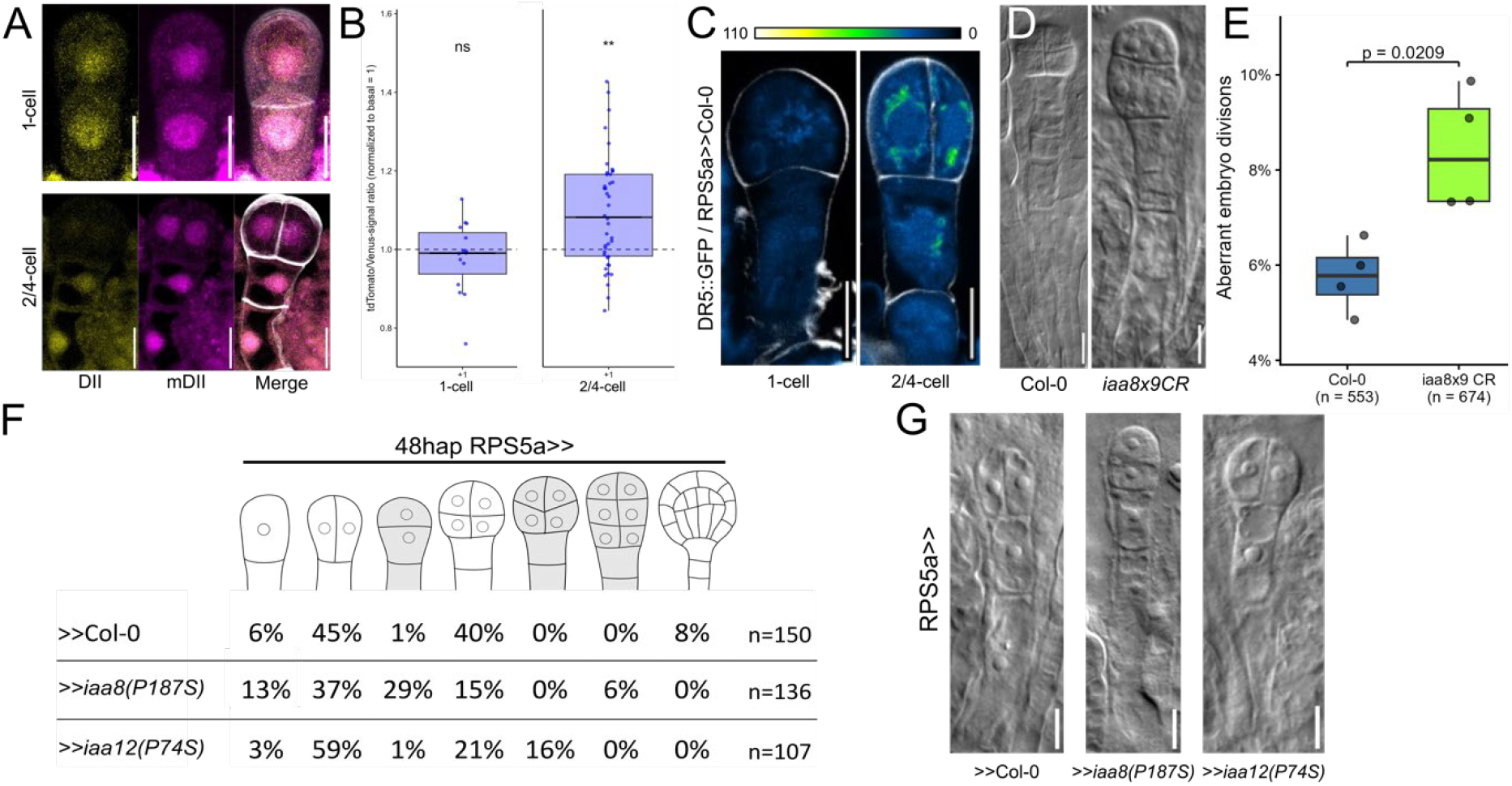
Auxin distribution and *IAA8* function in the early embryo. (A) DII-Venus (green) and mDIItdTomato (magenta) signals are shown as separate channels and as merged image with Renaissance SR2200 stain (white) for 1-cell and 2/4-cell embryos. Scale bar =10μm. (B) Calculated tdTomato/Venus signal ratio of the apical cell (position = +1), normalized to the most apical of suspensor cells (position = 0) of the embryo. Higher values indicate increased DII-Venus turnover and are interpreted as higher auxin levels. The population +1 was compared with population 0 within a specific embryo stage and analyzed by Wilcox rank-sum test. (C) Octant stage embryos of WT and *iaa8×9CR* (right). Aberrant division of the apical cell leads to deformation of the early embryo. Scale bar =10μm. (D) Quantification of aberrant divisions of the apical cell in Col-0 and *iaa8×9 CR* mutant. Embryos and aberrant divisions were counted in double-blind experiment. Statistical differences were determined using Wilcox rank-sum test. (F) *pDR5::ER-GFP* signal in early *pRPS5a>> Col-0* embryo. Scale bar = 10μm. (E) Quantification of embryo phenotype identification of F1 hybrids of *pRPS5A* activator line with different effector lines 48hap. (G) Comparison of 48hap F1 hybrid embryos. Scale bar 10μm.

As the known auxin response modules in the embryo at the 1-cell stage are already differentially expressed at this stage (Hamann *et al*., 2002; Rademacher *et al*., 2012) and its key players *MONOPTEROS* (*MP*) and *BODENLOS* (*BDL*) are expressed in an auxin-inducible way, additional auxin-independent cues are needed to initiate the embryonic patterning process. We rationalized that these might act on key components of the auxin response module at this stage to explain the differential *DR5rev::erGFP* activity in the absence of a detectable auxin gradient. Existing transcriptome data with spatial and temporal resolution during the first stages of embryonic development indicated that *IAA8* is expressed at high level at the 1-cell stage (Figure S1A; Zhao *et al*., 2019). IAA8 together with IAA9 and IAA27 belongs to a small subclade of the Aux/IAA protein family (Liscum and Reed, 2002) and *IAA9* transcripts can also be detected in early embryos at moderate levels (Figure 1A). We confirmed the early expression of *IAA8* by transcriptional fluorescent reporters and whole-mount RNA in-situ hybridization (Figure S1B-J). Both independent methods indicate that IAA8 is expressed in both cells at the 1-cell stage and their direct descendants. This is expected for a factor potentially involved in instructing differences in these cells. It also suggests that IAA8 expression is independent of transcriptional auxin responses as its expression pattern deviates from that of the auxin response reporter *DR5rev::erGFP*.

The turn of division plane orientation in the apical cell is one of the hallmarks of the initial embryonic auxin response (Hamann *et al*., 1999; Yoshida *et al*., 2014). If an auxin concentration gradient is only established at a later stage of embryonic development, we hypothesized that loss of the initial instructive factors should lead to a delay in embryo initiation before the 4-cell stage. We therefore analyzed division plane orientations in wild-type and *iaa8 iaa9* CRISPR (iaa8×9CR) double mutant embryos (Figure 1D, E and Figure S2A). We could detect a significant increase in atypical horizontal division planes that follow the minimal cell wall rule (Figure 1D, E), reminiscent of extra-embryonic development, emphasizing the importance of *IAA8* and *IAA9* in embryo initiation. It also highlights functional specificity of *IAA8* and *IAA9* in the process of initiating embryogenesis, as loss-of-function mutations in Aux/IAA genes barely show discernible phenotypes attributed to high genetic buffering in this multi-gene family (Abley *et al*., 2016; Overvoorde *et al*., 2005).

To determine whether *IAA8* indeed has a specialized role in early embryonic development, we expressed an auxin-insensitive, stabilized version of IAA8 (iaa8(P187S)) by transactivation. We included in this experiment IAA12/BDL (iaa12(P74S)/bdl; (Hamann *et al*., 2002)), a well-established factor of embryonic patterning in the globular embryo (Figure S2B, C). A GAL4>>UAS-based transactivation system was used to overcome embryo-lethality and to ensure similar expression of these genes using the ubiquitously active *RPS5A* promotor (Weijers *et al*., 2003). While patterning defects were obvious in *iaa12(P74S)* transactivated lines in the globular embryo which matched previous reports, early developmental defects at the 2/4-cell stage were detectable only at low frequency (Figure 1F, G). In transactivation lines with stabilized *iaa8(P187S)*, however, strong defects in 2/4-cell embryos were observable at a high frequency with apparent lack of embryo initiation (Figure 1F, G), indicating that *IAA8* can fulfill a more prominent function at this developmental stage. Expression of wild-type *IAA8* did not produce any discernible phenotype, confirming that the observed effect is not a consequence of mere overexpression (Table S1). As the resulting filament-like structures had the appearance of basal daughter cells without embryo (Figure 1G), we wanted to explore the molecular nature of these cells and analyzed the expression of known embryo-and early suspensor-specific genes by whole-mount RNA in situ hybridization. Expression of embryo-specific genes, such as *HANABA TANARU* (*HAN*; (Nawy *et al*., 2010)), *DORNRÖSCHEN* (*DRN*; (Chandler *et al*., 2007)), and *WUSCHEL-RELATED HOMEOBOX2* (*WOX2*; (Haecker *et al*., 2004)) was lost in some of these structures but also expanded or mislocalized in other similar structures, indicating that no stereotypic pattern of cell identities seems to have manifested yet. This is supported by a similar result for early basal cell lineage-specific genes the expression of which was either absent, mislocalized or expanded in a seemingly stochastic manner (Figure S3A-G). This strong effect on cell identity was not observed in transgenic lines expressing stabilized *iaa12(P74S)* (Figure S3A-G), suggesting that IAA8 plays an important and specific role in the early cell fate specification that differs from canonical Aux/IAAs such as IAA12.

To test if the specific effect of stabilized IAA8 protein can be explained by specific ARF interaction partners, we used domain swapping experiment with IAA8-IAA12 chimeras. ARF interaction is determined by the C-terminal PB1 domain of Aux/IAA proteins. However, the specific effect of IAA8 on early embryonic patterning was only detectable in constructs with the N-terminal part of IAA8, while the IAA12-IAA8 chimera with the PB1 domain of IAA8 evoked IAA12-like effects (Figure S3H, I).

One difference that we could identify between IAA8 and IAA12 was the presence of canonical MPK6 phosphorylation consensus sequences (Sorensson *et al*., 2012), situated in the N-terminal region, that are conserved between IAA8, IAA9, and IAA27 but not present in IAA12 (Figure S4). IAA8 has recently been described as target of MPK6-dependent phosphorylation in the context of heat stress response (Kim *et al*., 2024; Wang *et al*., 2025). We could confirm the reported protein-protein interaction between IAA8 and MPK6 in yeast 2-hybrid assays (Figure S5). Control by MAP kinase-mediated phosphorylation could therefore be important for the functional specificity of IAA8 in this process.

The embryonic ERECTA-YODA pathway has been established as a central player in establishing the initial cell identities in the early embryo (Dresselhaus and Jurgens, 2021; Ueda *et al*., 2017; Ueda *et al*., 2011; Wang *et al*., 2019; Wangler *et al*., 2026) and has been implicated in affecting auxin signaling (Rademacher *et al*., 2012). Furthermore, the embryonic phenotype of stabilized iaa8(P187S) resembles constitutively active variants of the MAP3K YODA, acting upstream of MPK6 (Lukowitz *et al*., 2004). The observed phosphorylation site motifs in IAA8 could therefore be a link between auxin signaling and this MPK6-dependent pathway. Recently, it has been shown in the context of heat stress response that MPK6-dependent phosphorylation can change protein turnover of Aux/IAAs and this has been demonstrated for IAA8 as well (Kim *et al*., 2022; Kim *et al*., 2024).

We therefore wondered if MPK6-dependent stabilization of IAA8 could explain the observed discrepancy between activation of ARF-dependent gene expression and the absence of detectable auxin accumulation in the apical cell. We tested protein turn-over of IAA8 variants including phospho-mimic (IAA8D) and phospho-dead (IAA8A) versions in protoplasts at increased auxin concentrations while inhibiting protein synthesis (Figure 2A). We measured significantly slower protein decay of IAA8 in the presence of auxin when potential phosphorylation sites were replaced by negatively charged amino acids. A similar effect could be observed in iaa8(P187S) variants that carry an amino acid exchange that interferes with TIR1 binding. Replacing potential phosphorylation sites with uncharged alanine did not stabilize the protein but rendered it unstable. This confirmed that MPK6 phosphorylation sites influence IAA8 stability in vivo as reported for heat stress (Kim *et al*., 2024; Wang *et al*., 2025). To understand whether the results obtained in protoplasts reflect the situation in the early embryo, we investigated IAA8 protein abundance in the early embryo in transgenic lines expressing YPet-fusions of IAA8 variants under the control of the early embryo-specific *S4* promoter (Slane *et al*., 2014). Although this promoter is uniformly active in all cells of the early embryo, similar to the *IAA8* promoter, we could only detect accumulation of IAA8 wild-type YPet-fusion protein in the basal cells of the embryo (Figure 2B). Phospho-mimic variants on the other hand accumulated in all cells of the early embryo (Figure 2C), while non-phosphorylated versions only reached very low levels in all cells (Figure 2D), comparable to the abundance of the wildtype IAA8-YPet variant in the apical cell. Stabilized iaa8(P187S)-YPet accumulated similarly to phospho-mimic variants (Figure 2E). These results indicate that IAA8 shows differential turn-over rates in the two daughter cells and this can be influenced by the phosphorylation status of the IAA8 protein. Previous work on transcriptional targets of the ERECTA-YODA pathway has established that MPK6 is preferentially active in the basal daughter cell and its descendants (Chen *et al*., 2025b; Ueda *et al*., 2017).

**Figure 2:**
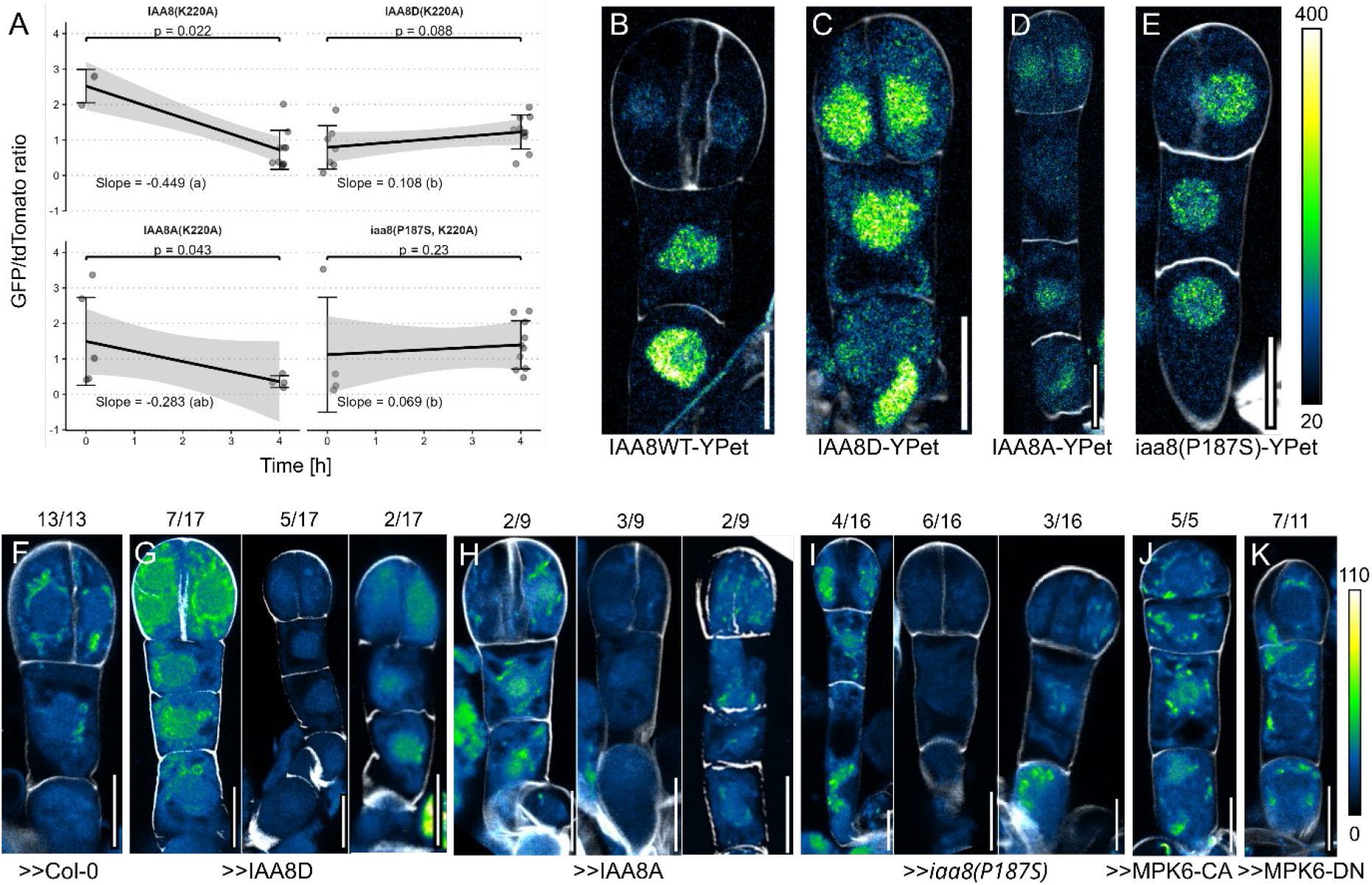
IAA8 turn-over and auxin responses in the early embryo. (A) Time course of protein turn-over of IAA8(K220A) variants in Arabidopsis protoplasts. Slopes: Interaction linear model with Multiple comparisons were adjusted using the Holm method. Compact letter indicates groups with non-significant differences in slope (α = 0.05). End point comparison: Wilcox rank-sum 48hap (B-E) Embryos expressing YPet fusion of IAA8-variants under the embryo-specific *S4* promoter. Cell walls are stained with Renaissance SR2200 to outline cells (white). YPet signal is shown in a green color scale (shown next to E). (B) Wild type IAA protein fusion (WT); (C) IAA8D, phospho-mimic variant; (D) IAA8A, phospho-dead variant; (E) iaa8(P187S), stabilized IAA8 variant. (F-K) F1 hybrid embryos showing *pDR5::ER-GFP* signal in different *UAS* effector lines crossed with *RPS5A:GAL4-VP16* driver lines. (F) Effector = Col-0 wildtype (>>Col-0). Ectopic DR5-signals can be detected in embryos expressing phospho-mimic IAA8 variants (G; >>IAA8D), phospho-dead IAA8 variants (H; >>IAA8A), stabilized IAA8 variants (I; >>iaa8(P187S)), or expressing constitutively active MPK6 versions (J; >>MPK6-CA), respectively. Frequencies of *DR5* activity patterns are given as numerical values above the image. The difference between the number of embryos with expanded, reduced, or inverted DR5 pattern and the total number given is accounted for by embryos that show *DR5* activity that cannot be distinguished from wild-type distribution. Scale bars = 10μm.

To determine whether IAA8 protein abundance has any effect on transcriptional auxin response, we monitored *DR5rev::erGFP* activity in the transgenic lines expressing different phospho-variants of IAA8. In wild type plants crossed with the *RPS5A* driver and *DR5rev::erGFP* reporter, we could observe weak GFP signal preferentially in apical cells (Figure 2F). When expressing IAA8A, IAA8D or iaa8(P187S) variants in all cells, we detected stochastic behavior of the GFP signal in all cells. For all three IAA8 variants, we observed GFP uniformly present in all cells, lost in all cells or preferentially apically or basally localized. This indicates that apical-basal domains are lost and cell identities are stochastic (Figure 2G-I). Without pre-patterning by differential IAA8 phosphorylation, auxin responses appear to be random, and no stable embryonic axis has been established.

Local auxin responses are critical for normal embryo development. We therefore analyzed the effect of changing phosphorylation status and accordingly IAA8 stability in the early embryo. Expression of wild-type IAA8 variants did not change embryonic patterning. However, when expressing IAA8A phospho-variants, we observed strong embryonic defects and malformed suspensors at the globular stage. Similarly, in variants that mimic phosphorylation, we detected club-like structures without clear embryo or suspensor formation, indicating that early cell fate decisions between embryonic and extra-embryonic development were disturbed (Figure S6B). Additionally, we observed a significant increase in aberrant division plane orientation in the apical cell at the 1-to 2-cell transition for both IAA8D and IAA8A variants, although less frequent than in iaa8(P187S) mutants (Table S1).

Taken together, these results support a scenario where ERECTA-YODA pathway activity leads to phosphorylation of IAA8 in the basal daughter cell, stabilizing it against auxin-dependent protein turnover. In the apical cell, non-phosphorylated IAA8 can be degraded independently of a steep auxin gradient or maximum in this cell.

According to this model, constitutive activity of the embryonic ERECTA-YODA pathway is predicted to suppress auxin responses in the early embryo, while loss of signaling activity would lead to ectopic auxin responses in the basal cell and the suspensor. To test this, we transactivated constitutively active MPK6-CA and dominant-negative MPK6 kinase-dead (MPK6-DN) variants under control of the ubiquitous *RPS5A* promoter. Expression of MPK6-CA resulted in filament-like structures similar to transactivation of stabilized *iaa8(P187S)* with uniform ectopic *DR5rev::erGFP* activity. Expression of MPK6-DN resulted in similar ectopic *DR5* activity while showing a milder morphological phenotype (Figure 2J, K), indicating that differential auxin responses are impaired in these lines. Correct patterning of the early embryo therefore relies on differential stabilization of IAA8 along the apical-basal axis.

Previous work reported that suspensor-specific expression of stabilized IAA12 variants led to callus-like proliferation of the suspensor and formation of suspensor-derived embryos (Rademacher et al., 2012). This seems to be in direct conflict with our model that relies on phosphorylation-dependent stabilization of IAA8 for normal embryonic patterning. We would therefore expect that in contrast to *IAA12*, suspensor-specific expression of stabilized *IAA8* would be tolerated by the plant. To test this and to confirm previous reports, we transactivated stabilized versions of IAA8 and IAA12 under control of the suspensor-specific *ARF13* promoter (Figure S6A). Confirming previous findings, suspensor-specific expression of stabilized *iaa12(P74S)* led to strong defects in suspensor development. In contrast, similar expression of stabilized *iaa8(P187S)* was tolerated by the embryo and only mild patterning defects could be detected after the globular stage likely due to prolonged, ectopic expression. These findings support the hypothesis that normal embryonic patterning involves specific stabilization of IAA8 in the early basal lineage.

To make sure that the observed phenotype in stabilized IAA8 variants is indeed connected to its canonical function in suppressing ARF activity, we targeted a range of A- and B-type ARFs (ARF2, 4, 5, 6, 7, 8) with an artificial micro-RNA (amiRNA). Expression of the amiRNA led to similar morphological changes as found in *iaa8(P187S)* mutants (Figure S6C), confirming that IAA8 likely acts in a canonical way in modulating ARF activity.

## Discussion

Auxin-dependent patterning has been described as a partially self-organizing system. A nice example for this is PIN1 in the shoot apical meristem, where its polarity is oriented towards an existing auxin maximum, thus increasing the auxin gradients steepness (Benkova *et al*., 2003; Huang *et al*., 2026). A similar scenario was envisioned for PIN7 in the early embryo, where it localizes in the basal cell towards the apical cell that shows an auxin response (Lau *et al*., 2012). Our data however shows that the first differential auxin response in the apical cell relies on MPK6-dependent, repressive Aux/IAA stabilization in the basal cell and that PIN7 activity only establishes an apical auxin maximum at a later stage. Differential Aux/IAA stabilization therefore precedes any auxin gradient and kick-starts the embryonic patterning process.

Changing auxin responsiveness by phosphorylation of Aux/IAA proteins seems to be an emerging theme as this protein family has recently been identified as downstream target of the auxin-sensitive receptor kinase TMK1 (Cao *et al*., 2019). And many other Aux/IAA proteins are reported to be phosphorylated (Kim *et al*., 2022; Kim *et al*., 2024; Noryang *et al*., 2026; Zheng *et al*., 2026). There is also evidence that Aux/IAAs are not the only component of the auxin transport and response machinery that is controlled by MAPK-dependent phosphorylation. The hydrophobic loop of PIN proteins is important for PIN activity and localization in a phosphorylation-dependent manner and PIN1 has been described as substrate for MPK6 (Huang *et al*., 2026; Janacek *et al*., 2024). Furthermore, phosphorylation of ARF7 and 19 modulates auxin responsiveness during lateral root formation (Cho *et al*., 2014). It will therefore be interesting to see if the first auxin response in the embryo is controlled by additional layers of MPK6-dependent regulation. For example, the differential expression of PIN7 and the stimulation of its activity would enhance the effect of MPK6-dependent IAA8 phosphorylation in defining apical-basal cell identities. The auxin flux generated by differential expression of *PIN7* in the basal cell will eventually lead to auxin accumulation in the apical cell, as seen by R2D2 measurement in 2/4-cell embryos. This might explain the rather mild defects observed in *iaa8 iaa9* double mutants. Future research needs to address if the *PIN7* gene and its product are targets of the embryonic YODA pathway.

Furthermore, in many patterning processes such as in the shoot apical meristems or during lateral root formation, auxin is instrumental (Abley *et al*., 2016; Bhatia and Heisler, 2018; Burian *et al*., 2026; Burian *et al*., 2022; Dubrovsky *et al*., 2008). But these processes are also known to be controlled by receptor kinase and MAP kinase signaling (Clark *et al*., 1997; De Smet *et al*., 2008; Hu *et al*., 2018; Oh *et al*., 2018; Roberts *et al*., 2016; Torii *et al*., 1996). It is therefore fascinating to speculate if auxin responses in these contexts could also be primed by preceding receptor signaling.

Previous reports established the effect of phosphorylation on IAA8 protein turn-over during heat-stress (Kim *et al*., 2024; Wang *et al*., 2025). However, the biological meaning in this context is hard to interpret. MPK3/6 signaling has been described in stress responses and innate immunity in addition to its role in plant development. Stabilizing IAA8 by phosphorylation during heat stress showed effects on MYB and bZIP transcription factor expression, which are involved in different contexts also ranging from stress responses to hormone signaling and development (Droge-Laser *et al*., 2018; Dubos *et al*., 2010). This raises the question whether IAA8 is an integral player of heat stress response or if its MAPK-dependent stabilization is an unavoidable cross talk between stress responses and developmental signals?

Recent reports indicate that the interaction of AUX/IAA proteins with TIR1 serves two purposes: Aux/IAA turnover and local cAMP production (Vanneste *et al*., 2025). Notably IAA8 phosphorylation, in contrast to stabilizing DII mutations (iaa8(P187S)), prevents ubiquitination but does not interfere with TIR1 binding (Kim *et al*., 2024). It will therefore be interesting to see in future studies if phosphorylated IAA8 can still trigger cAMP production in TIR1 complexes or whether phosphorylated IAA8 proteins bind to TIR1 in a non-functional way. According to proposed models, phosphorylated IAA8 bound to its target ARFs would still promote local cAMP production, stimulating ARF activity (Chen *et al*., 2025a). This would essentially override repression and induce an auxin response. However, this is not what we can observe by monitoring *DR5* activity, and auxin responses seem to be repressed in the basal cell by MPK6. Clearly, more research is necessary to characterize the interplay of these two novel mechanisms within the auxin response machinery.

*IAA8* belongs to a subclade with three members in Arabidopsis that is phylogenetically conserved (Liscum and Reed, 2002). Loss of *IAA8* and *IAA9* function impaired the initial embryo formation and delayed the patterning process. In many plant families, including those of grasses, the embryonic patterning process is not as strict as in *Brassicaceae* and seems to start at a later stage of embryogenesis. In rice, *OsWOX8/9A*, the homologue of *AtWOX8*, is under control or MPK6 and expressed in the basal region of the embryo, but auxin responses are detected only at later stages of development (Ishimoto *et al*., 2019). It will therefore be interesting to see if early embryonic development in the agriculturally important grass family relies on MPK6-dependent prepatterning or if the absence is one reason for the seemingly less organized embryonic patterning.

Auxin has often been described as a plant morphogen and has been attributed the ability to self-organize as seen during canalization (Abley *et al*., 2016; Benkova *et al*., 2003; Bennett *et al*., 2014; Bhatia and Heisler, 2018; Burian *et al*., 2026; Grieneisen *et al*., 2007; Lau *et al*., 2011). The observed influence of receptor signaling adds an additional layer of complexity and suggests that feedback loops in these self-organizing systems might be more complex than anticipated. The presented data imply that MPK6-dependent phosphorylation of Aux/IAA proteins forms a sensitivity difference, which in turn gives rise to a robust differential response at uniform auxin concentrations, essentially pre-patterning the embryo before auxin distribution matters.

## Materials and methods

### Plant materials and growth conditions

*Arabidopsis thaliana* strains used in this research were in the Colombia-0 (Col-0) background. All seeds were surface sterilized before germination. Plants were grown in long day conditions (16h light/8h dark) at 22°C and 50% humidity. T-DNA insertion lines *iaa8-1, iaa9-1* (Overvoorde *et al*., 2005) have been described previously. The R2D2 transgenic reporter line, the *pDR5rev::erGFP* line and the *ACT_RPS5A* activator line were published previously (Friml *et al*., 2003; Liao *et al*., 2015; Weijers *et al*., 2003). Similar activator lines that contain *pUAS::NLS-tdTomato*, resulting in *pRPS5a::VP16>>tdTomato* were constructed. *pUAS* represents five direct repeats of the Gal4 Upstream Activating Sequence (*UAS*) followed by a minimal 35s promotor (Engineer *et al*., 2005; Moore *et al*., 2006).

### Generating CRISPR lines

CRISPR-based loss-of-function alleles were essentially generated as previously described (Chen *et al*., 2025b). Primers for *IAA8* (AT2G22670) and *IAA9* (AT5G65670) locus (Table S2) were determined using CRISPR-P2.0 (Liu *et al*., 2017). sgRNAs were chosen to cut in the 5’ and 3’ UTR to achieve a complete removal of the coding region (Figure S2).

### Mutagenesis, cloning and transgenic lines

Plasmid vectors were constructed using the In-Fusion method (TAKARA BIO EUROPE S.A.S., Saint-Germain-en-Laye, France). The necessary primers were designed using the online tool provided by the manufacturer (https://www.takarabio.com/learning-centers/cloning/primer-design-and-other-tools).

Linear DNA fragments were amplified by PCR or generated by restriction enzyme-based linearization of plasmid vectors. Constructs for plant transformation were cloned into pBayBar or pBayHyg binary vectors(Wang *et al*., 2021). Additional constructs for plant transformation were cloned into pGreen II binary vectors (Hellens *et al*., 2000).

For genetic rescue constructs, the second intron was removed from the *IAA8* genomic sequence using IAA8i2del-F/R primers. This allows independent genotyping the genetic background in transgenic lines that contain a *IAA8* transgene. Bases Serine 33, Serine 80, Serine 91, Threonine 91 and Serine 152 were mutated using In-Fusion site directed mutagenesis. *iaa8(P187S)* mutant constructs were generated using IAA8(P187S)-F/R primers. MPK6 genomic sequences were mutated to create dominant negative (DN), and constitutively active (CA) variants. In the CA-variant coding regions were changed to result in Y144 to C transitions (Berriri *et al*., 2012). For dominant-negative variants, we replaced the TEY motif in the activation loop, amino acids 221-223, with an AEF amino acid sequence (Bush and Krysan, 2007).

*pS4::IAA8X-YPet:tUBQ10* was constructed by inserting the *IAA8* coding sequence into *pS4::YPet:tUBQ10* (Slane *et al*., 2014) using S4-IAA8-YPET-insert-F/R primers. *The pIAA8::GAL4-VP16-UAS::NLS-tdTomato:tIAA8* transcriptional reporter was constructed inserting 5000bp up-stream of the second ATG of the *IAA8* locus as well as 2500bp down-stream, after the *IAA8* STOP codon into a GAL4-VP16-UAS:tdTomato target vector. *pUAS::IAA8X:UBQ10* effector constructs were cloned by inserting variants of the *IAA8* CDS into the *pUAS::_:tUBQ10* effector vector using UAS-IAA8-intro-F/R primers. *pUAS::MPK6-CA, pUAS::MPK6-DN*, and *pUAS::iaa8(S187P)* were constructed by introducing the respective coding-regions into the pUAS::_:tNOS effector vector. For MPK6-CA and MPK6-DN the previously described mutated genomic sequences were inserted from the first ATG until STOP, 2217bp in total. For *iaa8(S187P)* the previously described mutated genomic sequence was inserted using the *IAA8*.*4* gene model, starting at the first ATG until 2238bp downstream of the STOP, 3821 bp in total.

Completed clones were confirmed using Sanger sequencing. The binary vectors were transformed into Agrobacterium tumefaciens strain GV3101, containing the *pSoup* plasmid. Plants were transformed by using floral dip. Positive transformants were selected on ½-strength Murashige Skoog medium, 0,8% Agar and PPT (15mg/l) or hygromycin (25mg/l). *p35s::IAA8X(K220A)-GFP:tCaMV* constructs used for protoplast transfections were generated using a *pJIT60* backbone (Lau *et al*., 2011). *IAA8(K220A)* variants were produced by site-directed mutagenesis using IAA8(K220A)-F/R primers to prevent forming of large protein oligomers in protoplasts. The *IAA8(K220A)* mutation containing PB1 domains fused to GFP were introduced into the pJIT60 target vector using pJIT-KtoA-GFP-F/R primers. Into the resulting vector, IAA8 sequences missing the PB1 domain sequence were introduced between the 35s promotor sequence and the mutated PB1 domain using pJIT-IAA8-Insert-II-F/R primers. *p35s::NLS-tdTomato:tCaMV* constructs were constructed by inserting the tdTomato sequence fused to a NLS sequence into the target vector between the 35s promotor sequence and CaMV terminator using the pJIT60-NLS-tdTom-F/R primers. Completed clones were confirmed using Sanger sequencing.

### Transactivation F1-hybrid analysis

For the analysis of F1-hybrids the pUAS/VP16-transactivation system was employed using stable effector and activator lines (Moore *et al*., 2006). Inflorescences of maternal crossing partners were emasculated and the exposed stigma pollinated after 24h. The fertilized siliques were grown on the plant for 48-72hap and harvested for the respective experiment.

### RNA in-situ hybridization

The probes were amplified directly from cDNA from seedlings except for *WOX8*, which was amplified from a plasmid instead, using the primers listed in Table S3. They were transcribed in vitro and labelled with up to a 1:3 ratio of UTP:Digoxigenin-11-UTP (Roche) using the T7 Polymerase (Thermo Scientific). Proper labelling was checked with a dot blot using DIG-labeled Control RNA (Roche) as reference. The UAS lines were emasculated and pollinated the next day with the driver line RPS5a. Two days after pollination the siliques were fixed, embedded and the in-situ hybridization was performed as described previously (Wang *et al*., 2021). The in-situ material was mounted in 15% glycerol.

### Microscopy

Siliques of the desired stage were fixed on double sided tape and cut open with tweezers. For confocal imaging, ovules were removed from the silique and immediately placed in a drop of 4% (w/v) paraformaldehyde in PBS (pH 6.9) solution with 0.1 to 0.3% of Renaissance 2200 stain (Musielak *et al*., 2015), depending on the embryo stage, on a microscopy slide as previously described (Musielak *et al*., 2016). The cover slide was placed on the ovules in solution and pressed on lightly with tweezers to rupture the ovules and release the embryos. *pS4::IAA8-YPet* variants were imaged at the Zeiss LSM880 inverse confocal microscope (Zen Black software; objectives 63x 1.2 W). Renaissance (Excitation 405nm, detection 414nm-450nm) and YPet (Excitation 514nm, detection 517nm-544nm) signal were recorded in Z-stacks and images analyzed using the FIJI software (Schindelin *et al*., 2012). *IAA8* promotor activity was imaged at the Zeiss LSM880 inverse confocal microscope (Zen Black software; objectives 63x 1.2 W). Renaissance (Excitation 405nm, detection 415nm-450nm) and tdTomato (Excitation 561nm, detection 570nm-597nm) signal were recorded in Z-stacks and images analyzed in FIJI software. R2D2 measurements were conducted at the LSM880 inverse confocal microscope (Zen Black software; objectives 63x 1.2 W). Renaissance (Excitation 405nm, detection 415nm-450nm), mVenus (Excitation 514nm, detection 517-545nm) and tdTomato (Excitation 561nm, detection 570nm-633nm) signals were recorded in Z-stacks on separate tracks to prevent signal bleed-through. *pDR5rev::erGFP* images were taken at the Leica SP8 upright confocal microscope (LAS X software, resonant scanner, 63x/1.2 W). Renaissance (Excitation 405nm, detection 414nm-450nm) and GFP (Excitation 488nm, detection 500-520nm) were recorded using the resonance scanner and very high averaging to prevent photobleaching.

For differential interference contrast (DIC) images ovules were removed from the replum and placed in Hoyer’s mounting solution (Chloral hydrate, water, and glycerol (ratio w/v/v: 8:3:1) and 10% gum arabic 2:1 mixture) or Chloral hydrate. DIC images were recorded using the Zeiss Axiophot (Zen software, AxioCam 512 color, Plan Neofluar 20x/0.5, 2x magnification), Zeiss Axio Observer (Zen software, AxioCam 305 color, AxioCam 820 mono, Plan-APOCHROMAT 40x/0.95 Korr) and Zeiss Axio Imager Z.1 (AxioVision software; AxioCam HR camera; Plan-APOCHROMAT ×40 and x10).

### Auxin measurements using the R2D2 reporter

The recorded R2D2 Z-stacks were analyzed in FIJI software. After maximum projection of the Z-stacks, gray values (GV) in the nucleus were measured for each channel. Embryos for 1-cell and 2/4-cell stage were analyzed. The recorded GVs were used to calculate the ratio of tdTomato signal:Venus signal and normalized to the most apical cell of the suspensor. The data was then processed and plotted using R Studio with Wilcox rank-sum test.

### IAA8 stability in protoplasts

Protoplasts were transfected using established protocols {Mehlhorn, 2018 #51400} with 20µl of plasmid transfection volume for 60µl Arabidopsis root-derived suspension cell culture protoplasts and incubated overnight. *p35s::IAA8(K220A)-GFP* variant plasmids were co-transfected with *p35s::NLS-tdTomato* plasmid. In single transfections, the total amount of plasmid was substituted with empty *pJIT60* plasmid, to ensure normalized expression of controls. After 24h in the dark, protoplasts were treated with 50µM IAA (Duchefa) and 50µM CHX (Duchefa) at t=0h. Protoplasts were placed on a slide with spacers and covered with a coverslip. Pictures were taken using the Zeiss Axio Observer (Axiocam 820 mono camera, 40x/0.95 objective). GFP (excitation 450-490 BP detection 500nm-550nm BP) and tdTomato (excitation 545nm-560nm BP detection 570nm-640nm BP) were recorded on separate tracks. Z-stacks of protoplasts were recorded at multiple time points and GV extracted using FIJI imaging software. Background GVs were measured and subtracted. Cross-channel signal was calculated and subtracted. GFP expression was normalized to the co-transformed *p35s::NLS-tdTomato* and non-co-transfected protoplasts filtered from the dataset to ensure analysis of protoplasts expressing both proteins.

### Embryo counting

To assess prevalence of phenotypes in embryos the hallmark phenotype of the turned apical division plane was chosen. Embryos were analyzed at the 2/4-cell transition stage and classified by the orientation of the division plane. For analysis the cleared embryos were counted using DIC microscopy. Embryos were counted according to the observable status of the apical cell, sorted for undivided, 2/4-cell correct division, 2/4-cell aberrant division and 8-cell or later. Counting was conducted in a double-blind setting to minimize observer bias influencing the assessment of phenotypes. After counting, data sets were pooled for replicates, thus single datapoints account for populations in an experiment rather than individuals. Data was assessed using R.

### Yeast experiments

The Split Ubiquitin-based protein interaction system was used as previously described {Asseck, 2018 #45808}. Bait vectors and prey vectors were transformed into THY AP4 and THY AP5 yeast strains respectively using YEAST Transformation kit (Sigma-Aldrich). The transformed yeast cells were plated on synthetic complete (SC) drop-out plates, leucine for bait and tryptophan and uracil for prey. Few colonies were inoculated into a liquid culture and grown overnight at 30°C with shaking. 1ml of the culture was pelleted and resuspended in 200µl of YPD. 20µl each of bait and prey were mixed in a 96 well plate according to required combinations and plated on YPD plates. After 8 hours the colonies were replica plated on SC drop-out plates (-adenine - histidine). To observe the interaction between bait and prey the colonies were plated on minimal medium with varying concentration of methionine and along with the dilutions (1/10, 1/100). To transform a library into yeast cells, we used yeast transformation kit from ClontechTM. In a screen where yeast cells contain bait + bridge + prey the minimal media was used. When only prey library was transformed into yeast cells bait, uracil synthetic complete drop-out plates were used.

### Data analysis

All data analysis was conducted using R Studio (R version 4.4.2, http://www.r-project.org)

## Acknowledgements

This project was funded by the Deutsche Forschungsgemeinschaft (DFG; German Research Council), grant number BA3356/3-1 (494821461) and the Max Planck Society.

We would like to thank Ole Herud, Steffen Lau and Ive De Smet for discussions during the early stage of the project, Chiara Leuzzi, Lara Kirchmair, Patricia Weber, Richard Gavidia, Steffi Zimmerman (ZMBP Tübingen) and Martin Vogt (MPI Tübingen) for technical assistance and discussions of the experiments, The ZMBP green house facility, ZMBP microscopy unit and the ZMBP transformation facility. We also would like to thank the Nottingham Arabidopsis Stock Centre (NASC) for providing sequence-indexed Arabidopsis T-DNA insertion mutants.

