## Supplementary figures and tables for "Before auxin distribution matters: Prepatterning of the Arabidopsis embryo axis by MPK6-dependent AUX/IAA8 phosphorylation"

**Supplementary files and Tables**

**Figure S1**: *IAA8* expression in the early embryo

**Figure S2**: CRISPR-based loss-of-function alleles of *IAA8* and *IAA9*, degron sequences and gene models

**Figure S3**: RNA *in situ* hybridization of apical and basal marker genes in RPS5A transactivation F1 hybrid embryos 48 hap.

**Figure S4**: Alignment of Aux/IAA protein sequences for IAA8, IAA9 and IAA27.

**Figure S5**: Protein-protein interaction assay using the MAP3K YDA as bait and MPK6 as untagged bridging protein

**Figure S6**: Embryonic phenotypes of transactivated embryos

**Table S1**: Quantification of transactivation F1 hybrid embryos 48 hap.

**Table S2**: Primer List,

**Table S3**: Primer for RNA *in situ* hybridization probes

**Figure S1:**

A

| Gene | EC | Zy14 | Zy24 | 1C | 32C |
|---|---|---|---|---|---|
| IAA12 (BDL) | 3.30 | 4.80 | 0.28 | 20.70 | 44.90 |
| IAA8 | 21.50 | 37.30 | 113.60 | 131.40 | 406.60 |
| IAA9 | 127.20 | 121.90 | 31.60 | 184.10 | 277.60 |
| PAP2 (IAA27) | 0.00 | 0.00 | 3.65 | 0.01 | 13.51 |
| PIN7 | 0.00 | 0.14 | 0.01 | 5.68 | 35.74 |

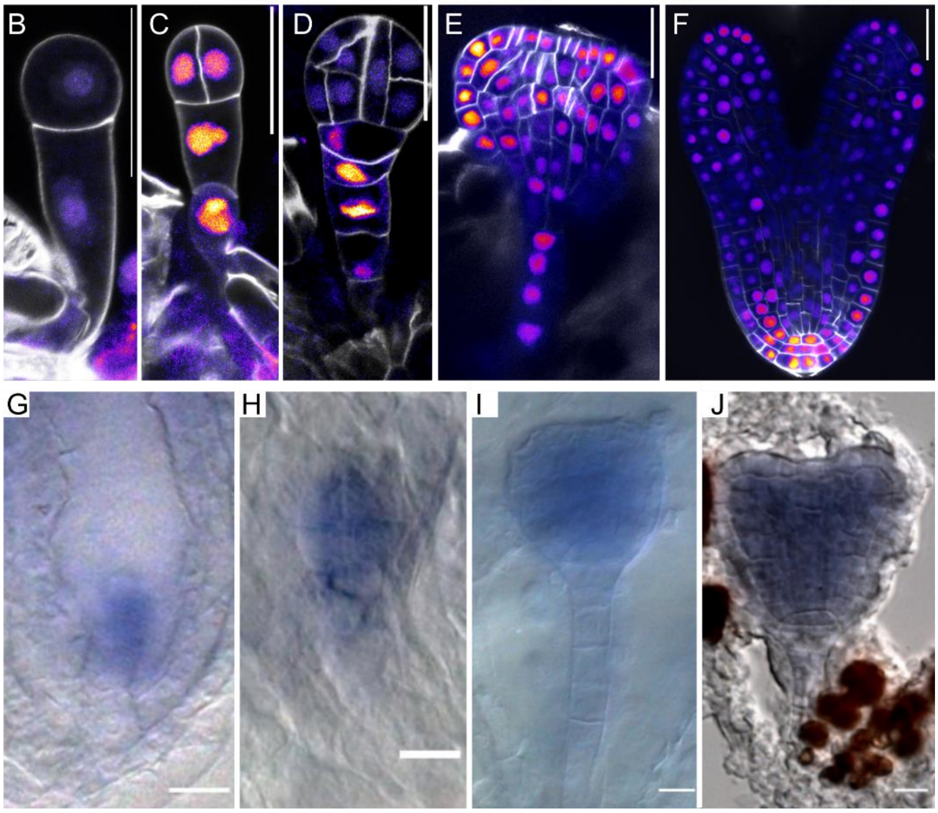


**Figure S1: *IAA8* expression in the early embryo**

(A) Expression of selected Aux/IAA genes and PIN7 in the early embryo according to publicly available transcriptome data (Zhao *et al*., 2019). EC, egg cell; Zy14, early zygote; Zy24, late zygote; 1C, 1-cell embryo after the first zygotic division; 32C, globular stage embryo.

(B-F) *IAA8* promoter activity in embryo stages from 1-cell (B), 2/4-Cell (C), dermatogen (D), transition stage (E), and Torpedo stage (F). Scale bar = 20um.

(G-J) in situ mRNA hybridization with *IAA8* antisense probe in early embryo stages, 1-cell (G), octant (H), transition stage (I) and early heart (J). Scale bar = 10um.

**Figure S2:**

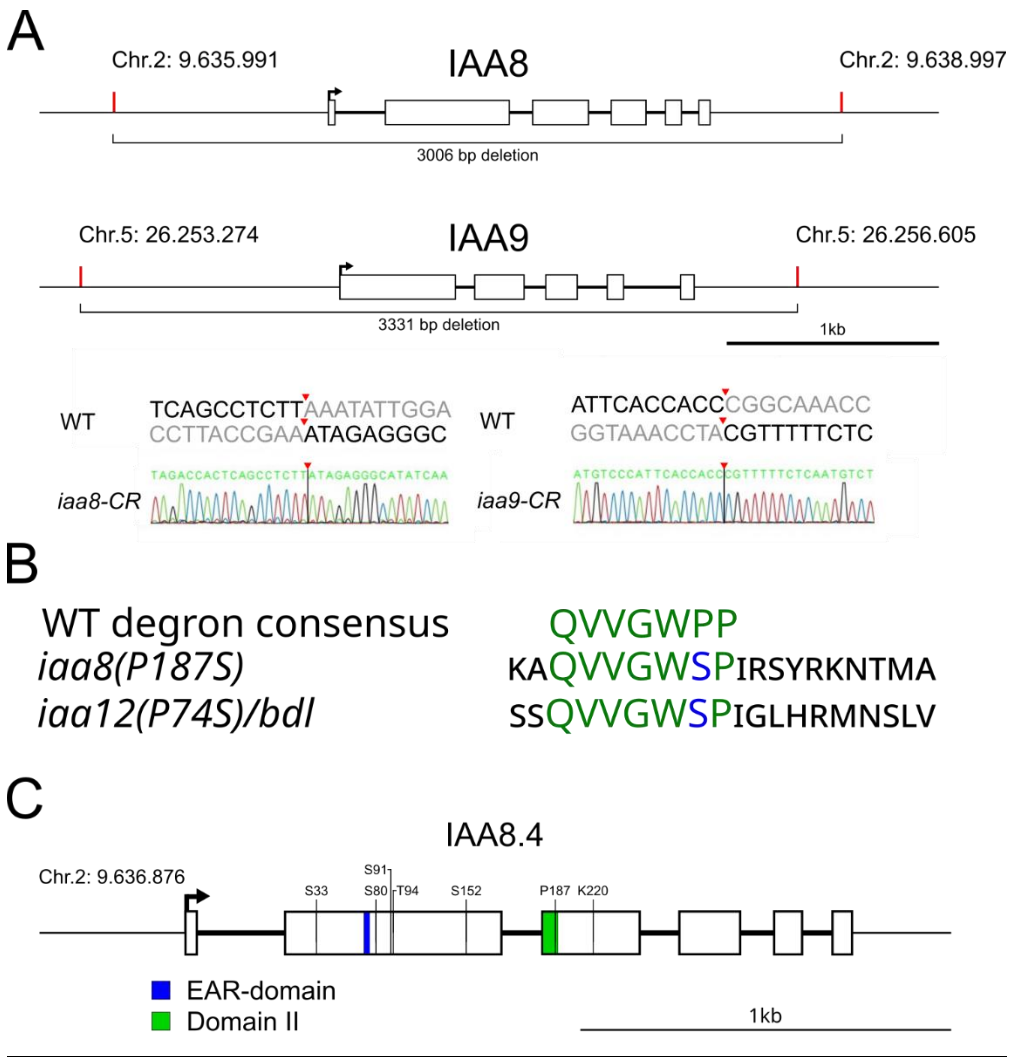


**Figure S2: CRISPR-based loss-of-function alleles of *IAA8* and *IAA9*, degron sequences and gene models**

(A) *IAA8* and *IAA9* loci marked with CRISPR sgRNA target sites (red). Positions annotated on Chromosomes (Chr.) on positive strand. Sanger sequencing results confirming complete removal of the locus. Red arrow heads and bars mark sgRNA target sites.
(B) Degron sequences of select Aux/IAAs, including surrounding amino acids. Degron domains are marked in green. The proline-to-serine exchange that stabilizes Aux/IAA proteins is marked in blue (Hamann et al., 2002).
(C) Gene model of *IAA8.4* (AT2G22670.4), which starts at the first ATG. Sites for modifications are marked, as well as EAR and DII domains. Position annotated on Chromosome (Chr.) 2 on positive strand. Size marker indicates 1 kb.

**Figure S3:**

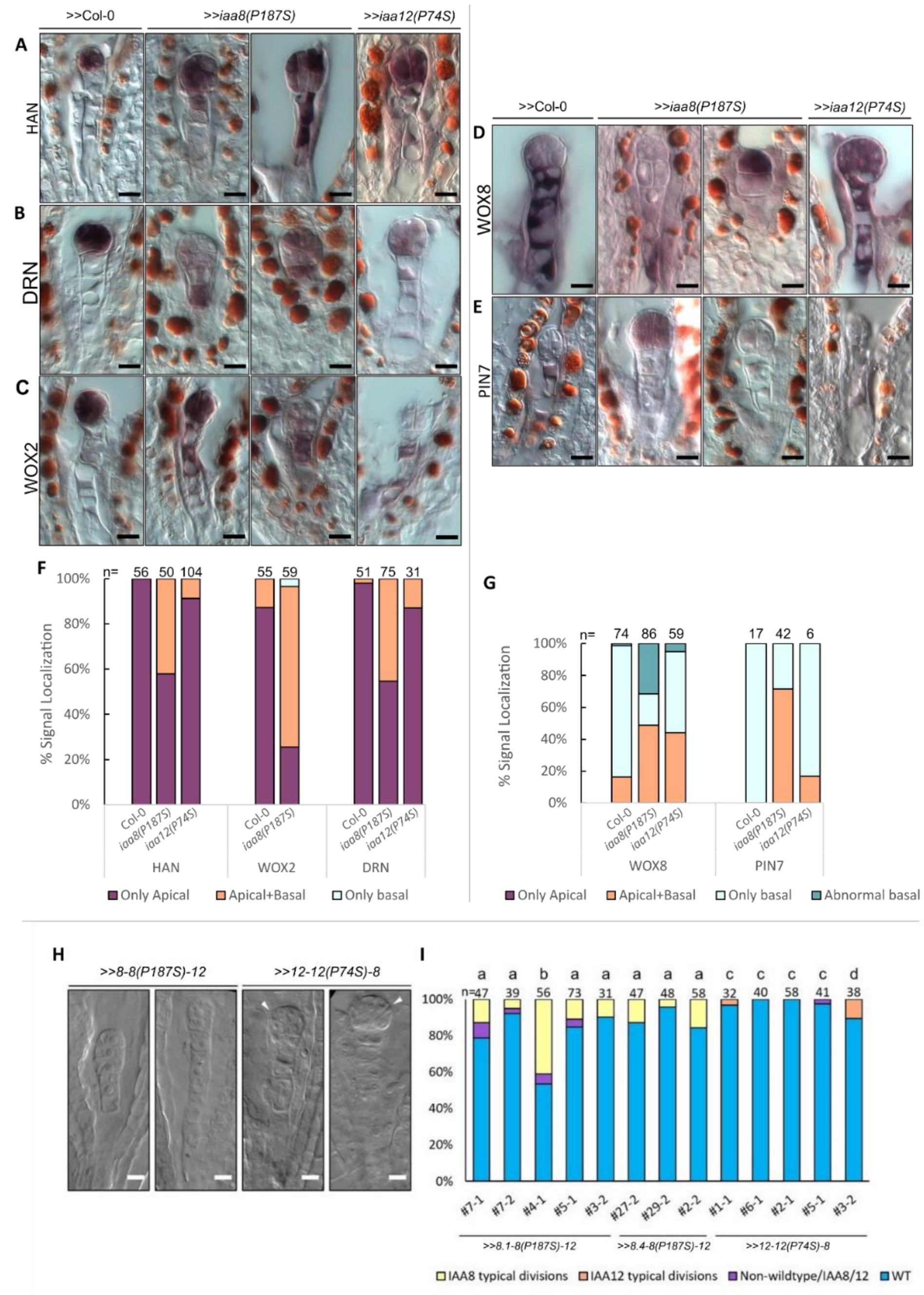


**Figure S3: RNA *in situ* hybridization of apical and basal marker genes in RPS5A transactivation F1 hybrid embryos 48hap.**

(A-C, F) DIC images (A-C) and quantification (F) of apical, embryonic marker genes, *HAN* (A), *DRN* (B), *WOX2* (C), loose their stereotypic pattern in stabilized *iaa8* variants.
(D-E, G) DIC images (D-E) and quantification (G) of basal marker genes, *WOX8* (D), *PIN7* (E), similarly seem to be stochastically expressed in stabilized *iaa8* variants.
(H, I) DIC images (H) and quantification (I) of phenotypic effects of chimeric *iaa8* and *iaa12* domain-swap variants. The IAA8-specific, aberrant cell divisions in the apical cell lineage are only detectable at high frequency if the N-terminal part is IAA8-derived.

**Figure S4:**

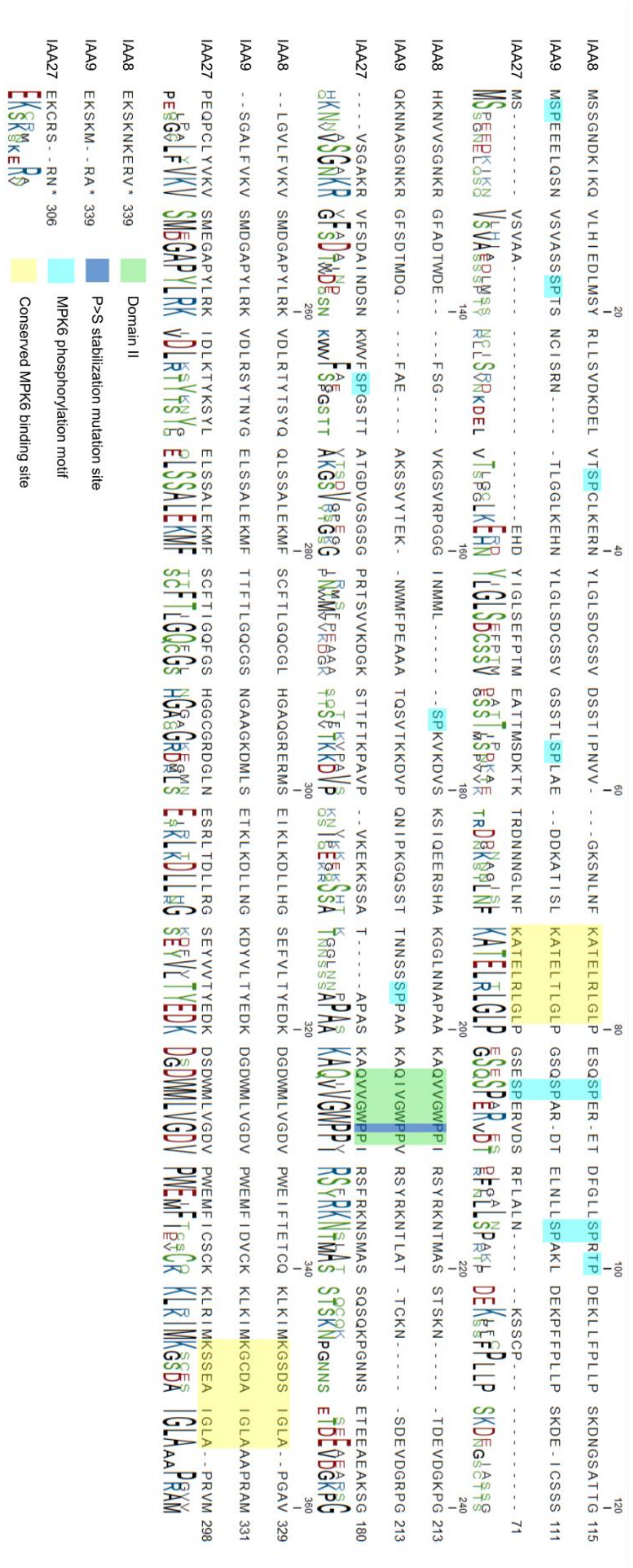


**Figure S4 Alignment of Aux/IAA protein sequences for IAA8, IAA9 and IAA27.** Regions of interest (potential MPK6 binding sites and phosphorylation sites, DII degron and conserved proline) are marked in color according to the legend.

**Figure S5:**

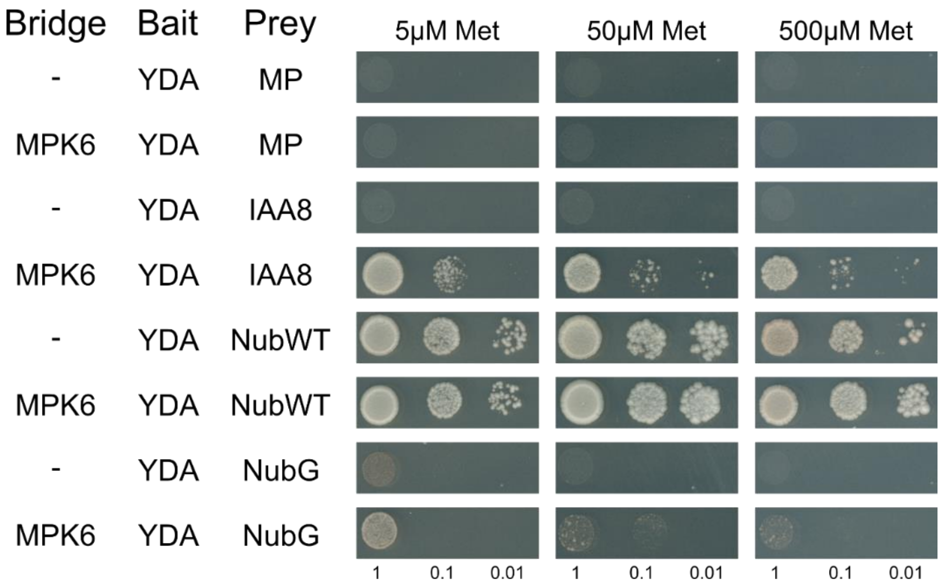


**Figure S5: Protein-protein interaction assay using the MAP3K YDA as bait and MPK6 as untagged bridging protein**

Yeast SUS bridge assay on SC drop-out plates with different Met concentrations (5µM, 50µM, 500µM). Increased Methionine concentrations lead to decreased prey expression to prevent auto activation. OD600 cultures were spotted as dilutions of 1x, 0.1x, 0.01x. NubWT is used as positive control, NubG as negative control.

**Figure S6:**

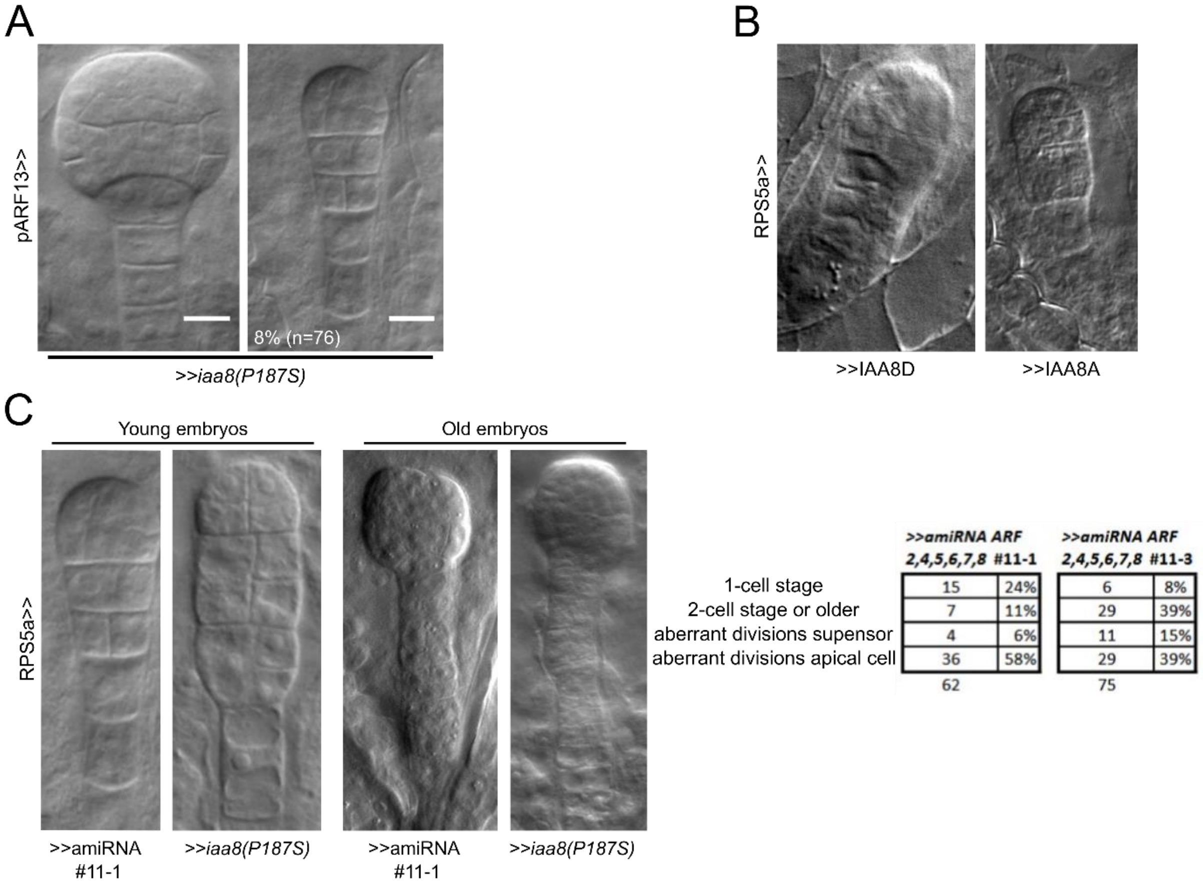


| | >>amiRNA ARF 2,4,5,6,7,8 #11-1 | | >>amiRNA ARF 2,4,5,6,7,8 #11-3 | |
|---|---|---|---|---|
| 1-cell stage | 15 | 24% | 6 | 8% |
| 2-cell stage or older | 7 | 11% | 29 | 39% |
| aberrant divisions supensor | 4 | 6% | 11 | 15% |
| aberrant divisions apical cell | 36 | 58% | 29 | 39% |
| | 62 | | 75 | |

**Figure S6: Embryonic phenotypes of transactivated embryos**

(A) F1 Hybrid embryos expressing >>*iaa8(P187S)* under the suspensor specific *ARF13* promotor (72 hap) display wild-type morphology. Weak defects can be observed at low frequency (right panel). No filamentous structure can be detected which are common with ubiquitous *iaa8(P187S)* expression (see figure 1). Scale bar = 10 um.

(B) F1 Hybrid embryos expressing *IAA8* phospho-variants under the *RPS5A* promotor (72 hap). Strong aberrant embryonic divisions are visible.

(C) Comparison of F1 hybrid mutants expressing either artificial miRNA against multiple ARFs (ARF 2, 4, 5, 6, 8) or stabilized *iaa8(P187S)* under the *RPS5A* promotor. Phenotypes at younger stages (left) and older stages (right) show resemblance in aberrant divisions of the apical tier and basal tier. Quantification of F1 hybrid lines expressing amiRNA against multiple ARFs (ARF 2, 4, 5, 6, 8) at 48 hap given as total number and percentage.

**Table S1:**

| Genotype | Frequency turned division planes | n= | p_value | significant |
|---|---|---|---|---|
| Col-0 | 1.63% | 123 | | |
| IAA8 (WT) | 1.60% | 188 | 0.98 | No |
| IAA8A | 9.74% | 154 | 0.00521 | Yes |
| IAA8D | 7.36% | 163 | 0.02619 | Yes |
| iaa8(P187S) | 18.02% | 172 | 0.00001 | Yes |

**Table S1: Quantification of transactivation F1 hybrid embryos 48 hap**. Frequency of turned apical cell divisions in % and total number (n) of analyzed embryos. Statistical significance of difference to RPS5a>>Col-0 embryos (Col-0). Two-proportion z-test $p<0.05$.

**Table S2: Primer List**,

Mutations marked in **bold lower case,** *guideRNA sequences in italics*

| | | |
|---|---|---|
| **IAA8i2del** | F | GCTGCCAAGGCACAGGTTGTTGGTTGGC |
| | R | CTGTGCCTTGGCAGCTGGTGCATTGTTC |
| **IAA8-P187S** | F | TTGGTTGG**t**CTCCAATCAGATCATACCGGAAG |
| | R | TTGGAG**a**CCAACCAACAACCTGTGCC |
| **IAA8-S33D** | F | AAGGATGAACTGGTTACG**gac**CCTTGTTTGAAAGAACGT |
| | R | ACGTTCTTTCAAACAAGG**gtc**CGTAACCAGTTCATCCTT |
| **IAA8-S80D** | F | CTTCCTGAGTCTCAA**ga**TCCTGAGAGAGAGACT |
| | R | AGTCTCTCTCTCAGGA**tc**TTGAGACTCAGGAAG |
| **IAA8-S91T94D** | F | ACTGATTTCGGTTTGCTG**ga**TCCGAGA**gat**CCCGATGAGAAGCTTCTC |
| | R | GAGAAGCTTCTCATCGGG**atc**TCTCGGA**tc**CAGCAAACCGAAATCAGT |
| **IAA8-S152D** | F | ATCAACATGATGTTG**gat**CCGAAAGTTAAGGAT |
| | R | ATCCTTAACTTTCGG**atc**CAACATCATGTTGAT |
| **IAA8-S33A** | F | AACTGGTTACG**g**CACCTTGTTTGA |
| | R | TCAAACAAGGTG**c**CGTAACCAGTT |
| **IAA8-S80A** | F | CCTGAGTCTCAA**g**CTCCTGAGAGA |
| | R | TCTCTCAGGAG**c**TTGAGACTCAGG |
| **IAA8-S91T94A** | F | CTG**g**C**t**CCGAGA**g**CACCCGATGAGAAGCTTCT |
| | R | TG**c**TCTCGG**a**G**c**CAGCAAACCGAAATCAGTCTCTCT |
| **IAA8-S152A** | F | AACATGATGTTG**g**CGCCGAAAGTT |
| | R | AACTTTCGGCG**c**CAACATCATGTT |
| **IAA8K220A** | F | GTTTGTG**gc**GGTGAGCATGGATGGTGC |
| | R | CTCACC**gc**CACAAACAGAACACCAAGACCAGG |
| **S4-IAA8-YPET-insert** | F | TTCCGGCAGGCTCGAGATGAGTTCTGGGAACGATAAGG |
| | R | TTTAGACACCATCCCGGGAACCCGCTCTTTGTTCTTCG |
| **pJIT60-NLS-tdTom** | F | GCAGGTCGACGGATCCATGGCTCCAAAGAAGAAGAGAAAGG |
| | R | TCAGCGTACCGAATTCTTACTTGTACAGCTCGTCCATGCC |
| **pJIT-KtoA-GFP** | F | GCAGGTCGACGGATCCGTTCTGTTTGTGGCGGTGAG |
| | R | TCAGCGTACCGAATTCTTACTTGTACAGCTCGTCCATGCC |
| **pJIT-IAA8-Insert-II** | F | CAGCCCAAGCTTGGCTGCAGATGAGTTCTGGGAACGATAAGG |
| | R | GCCACAAACAGAACACCAAGACCAGGTTTCCC |
| **UAS-IAA8-intro** | F | AACCTCTAGAACTAGTATGAGTTCTGGGAACGATAAGA |
| | R | TTGAACGATCCTCGAGTCAAACCCGCTCTTTGTTCT |
| **sgIAA8-A-CR** | F | *CCTCACCTTACCGAAATAGA*GTTTTAGAGCTATGC |
| | R | *TCTATTTCGGTAAGGTGAGG*CAATCACTACTTCGA |
| **sgIAA8-B-CR** | F | *GGGACCATCCAATATTTAAG*GTTTTAGAGCTATGC |
| | R | *CTTAAATATTGGATGGTCCC*CAATCACTACTTCGA |
| **sgIAA9-A-CR** | F | *TTCCAAGTCGGTTTGCCGGG*GTTTTAGAGCTATGC |
| | R | *CCCGGCAAACCGACTTGGAA*CAATCACTACTTCGA |
| **sgIAA9-B-CR** | F | *ATAGACATTGAGAAAAACGT*GTTTTAGAGCTATGC |
| | R | *ACGTTTTTCTCAATGTCTAT*CAATCACTACTTCGA |
| **MPK6 Y144C** | F | CAACGATGTTTACATCGCGTgTGAGTTAATGGACACTGATC |
| | R | GATCAGTGTCCATTAACTCAcACGCGATGTAAACATCGTTG |

**Table S3: Primer for RNA *in situ* hybridization probes**

| Probe | Primer pair 5´->3´ |
|---|---|
| ***IAA8*** | GCTTTTGTTAGCTTTCTCTGG<br>TAATACGACTCACTATAGGGTCAAACCCGCTCTTTG |
| ***PR2*** | AACAGACACAAAAGCATATAAGCA<br>TAATACGACTCACTATAGGGTATAAAAAAACAATATTAATTAG |
| ***PR8*** | TCTTACATTTTCAGCTGCT<br>TAATACGACTCACTATAGGGTGGTGAAAACGTGAATCTAAAAT |
| ***WOX2*** | CATGCAAACCATCGTCTTAAAACC<br>TAATACGACTCACTATAGGGCATAAAATTTATAATTTCATTAAACCTTCG |
| ***WOX8*** | TACACCATCATCATGTCCTCCT<br>TAATACGACTCACTATAGGGTATCCATAGCACCATAACATTTGC |
| ***DRN*** | TTTTTCCAACAAGAAATCTTCGCC<br>TAATACGACTCACTATAGGGCCAACATTGGGAAAGGTAGCAAC |
| ***PIN7*** | CGCTTCTTCTCTTTCTTTTCG<br>TAATACGACTCACTATAGGGTCCACAGCAGAGCTAAACCC |
| ***HAN*** | TACACTTAGGGTTTTCAAACCAG<br>TAATACGACTCACTATAGGGCTGGTGCTTCTTCTTGCATCT |
| | GGTTCAAACCGACCAGTACG<br>TAATACGACTCACTATAGGGCGAGAGACGATGTTGATGTTTAT |
| ***MP*** | CATCCGGAACCAAAGAAAGAAACTGTG<br>TAATACGACTCACTATAGGGTCTTGTACCTGACTGGTCTTTCAACAGC |
| | AGTTCGAGATCTTCATGAGAATACTTGG<br>TAATACGACTCACTATAGGGGAATCAGGAACACGTATAAGTGGC |
| | ACTCAGTTGAACGGTCTC<br>TAATACGACTCACTATAGGGTCTCCCGACTGACCCGGTT |
| ***GFP*** | CCACCTACGGCAAGCTGAC<br>TAATACGACTCACTATAGGGCCAGGATGTTGCC |
